# Neuromotor impairment, hearing loss and blindness in a preclinical mouse model of Charcot Marie-Tooth disorder

**DOI:** 10.1101/2020.06.02.130625

**Authors:** Sergio Gonzalez-Gonzalez, Chantal Cazevieille

## Abstract

Schwann cells produce myelin sheath around peripheral nerve axons. Myelination is critical for rapid propagation of action potentials, as illustrated by the large number of acquired and hereditary peripheral neuropathies, such as diabetic neuropathy or Charcot-Marie-Tooth (CMT) diseases, that are commonly associated with a process of demyelination. Peripheral neuropathy is a major complication of diabetes, and the pathomechanism of the disease remains poorly studied. Here, we studied the progressive demyelinating process, hearing impairment and blindness observed in the CMT1A mouse model C3. Our results confirm that these mice represent a robust and validated model to study the peripheral neuropathy induced by CMT disorder allowing to determine the efficacy of new pharmacological candidates targeting demyelinating diseases such as CMT1A disorder.

## Introduction

In the peripheral nervous system (PNS), Schwann cells (SCs) are responsible for myelin production, which contributes to axonal protection and allows for efficient action potential transmission (1, 2). Unfortunately, acquired and hereditary demyelinating diseases of the PNS are numerous and affect an increasing number of people (3).

Acquired demyelinating diseases are the most common, as they include diabetic peripheral neuropathy (4, 5), drug-related peripheral neuropathies, leprosy, and inflammatory neuropathies (6). Diabetic peripheral neuropathy is a major complication of diabetes type 1 and 2 and a cause of considerable morbidity (7, 8). Indeed, it has been reported that at least 50% of diabetic patients develop one or several forms of diabetic neuropathy within 25 years after diagnosis (7). This neuropathy affects both myelinated SCs and peripheral axons/neurons, leading to changes in nerve conduction, and it is often associated with demyelination in the long term (9). However, the physiological and molecular mechanisms that lead to these nerve defects remain unclear.

The etiologies of acquired and hereditary peripheral nerve diseases are diverse, but they all result in demyelination and subsequent neuronal death. Thus, an important challenge is to understand the cellular and molecular events that underlie demyelination of SCs.

Charcot-Marie-Tooth disease (CMT) is a hereditary motor and sensory neuropathy of the peripheral nervous system characterized by a progressive loss of muscle tissue and a dysfunction of the tactile sensation in different parts of the body (10). Currently incurable, this disease is the most prevalent hereditary neurological disorder and affects approximately one in 2,500 people. The most common type of CMT is CMT1A, characterized by a duplication of the pmp22 gene leading to an accumulation of the pmp22 protein in the Schwann cell and progressive demyelination (10, 11). PMP22 is a tetraspan glycoprotein contained in compact myelin of the peripheral nervous system. Duplication of PMP22 has been associated with the onset of Charcot-Marie-Tooth disease type 1A (CMT1A) (12).

The C3-PMP22 transgenic mice express three to four copies of a wild-type human peripheral myelin protein 22 (PMP22) gene (13). These mice present an age-dependent demyelinating neuropathy characterized by predominantly distal loss of strength and sensation. C3-PMP mice show no overt clinical signs at 3 weeks and develop mild neuromuscular impairment in an age-dependent manner. They have stable, low nerve conduction velocities similar to adults with human CMT1A. Myelination is delayed in these mice, and they contain reduced numbers of myelinated fibers at 3 weeks of age. There is no detectable loss of myelinated fibers in adult C3-PMP22 mice.

## Materials and methods

### Animal housing

Male CMT mice [B6.Cg-Tg(PMP22)C3Fbas/J, Jackson Labs], their controls [Non-carrier for Tg(PMP22)C3Fbas, Jackson Labs] were kept in the A1 animal house facility. Mice were housed in ventilated and clear plastic boxes and subjected to standard light cycles (12 hours in 90-lux light, 12 hours in the dark). Experiments started at 1 months of age (baseline) and finished at 6 months old. All animal experiments were approved by the Comité d’Ethique en Expérimentation Animale du Languedoc-Roussillon, Montpellier, France, and the Ministère de l’Enseignement Supérieur et de la Recherche, Paris, France

### Balance beam test

Balance beam test is a narrow “walking bridge” that rodents can cross to test balance and neurosensory coordination. The beam (thickness 6 cm) was elevated with the help of two feet with platforms to hold mice. The time required to cross the beam from side to side was quantified for each mouse. Each animal underwent 3 trials a day at 5minutes intervals. For each day, values from the 3 trials were averaged for each animal, normalized according to animal weight, and then averaged for each treated group.

### Rotarod

A rotating rod apparatus was used to measure neuromuscular coordination and balance. Mice were first given a 2-day pretraining trial to familiarize them with the rotating rod. Latency to fall was measured at a successively increased speed from 4 to 40 rpm over a 300-second time period. Each animal underwent 3 trials a day. For each day, values from the 3 trials are averaged for each animal, normalized according to animal weight, and then averaged for each group.

### Kondziela’s inverted screen test

The Kondziela screen test is a vertically positioned grid box, which allows the mouse to grab on the grids as it climbs down. This test allows to determine muscular strength and proprioception. The grip latency to fall was quantified for each mouse. Each animal underwent 3 trials a day at 5 minutes intervals. For each day, values from the 3 trials were averaged for each animal, normalized according to animal weight, and then averaged for each treated group.

### Sciatic nerve electrophysiology

Standard electromyography was performed on mice anesthetized with ketamine/xylazine mixture. A pair of steel needle electrodes (AD Instruments. MLA1302) was placed subcutaneously along the nerve at the sciatic notch (proximal stimulation). A second pair of electrodes was placed along the tibial nerve above the ankle (distal stimulation). Supramaximal square-wave pulses, lasting 10 ms at 1 mA for mice were delivered using a PowerLab 26T (AD Instruments). CMAP was recorded from the intrinsic foot muscles using steel electrodes. Both amplitudes and latencies of CMAP were determined. The distance between the 2 sites of stimulation was measured alongside the skin surface with fully extended legs, and NCVs were calculated automatically from sciatic nerve latency measurements. Only the left sciatic nerves were analyzed in this study.

### Sciatic nerve and auditory nerve TEM

The sciatic and auditory nerves of all mice were fixed for 20 minutes in situ with 4 % PFA and 2.5% glutaraldehyde in 0.1 M phosphate buffer (pH 7.3). Then, nerves were removed and post-fixed overnight in the same buffer. After washing for 30 minutes in 0.2 M PBS buffer, the nerves were incubated with 2% osmic acid in 0.1 M phosphate buffer for 90 minutes at room temperature. Then, samples were washed in 0.2 M PBS buffer, dehydrated using ethanol gradient solutions, and embedded in epoxy resin. Ultrathin (70-nm) cross sections were cut and stained with 1% uranyl acetate solution and lead citrate and analyzed using a HITACHI H7100 electron microscope at the COMET platform. Only the left sciatic nerves were analyzed in this study.

### Auditory brainstem responses (ABR)

ABRs are electric potentials recorded from scalp electrodes, and the first ABR wave represents the summed activity of the auditory nerve fibers contacting the inner hair cells. For ABR studies, mice were anesthetized using ketamine/xylazine mixture and body temperature is regulated using a heatingpad at 37 °C. Then, earphones were placed in the left ear of each mouse, an active electrode was placed in the vertex of the skull, a reference electrode under the skin of the mastoid bone and a ground electrode was placed in the neck skin. The stimuli consisted of tone pips of five frequencies (4 kHz, 8 kHz, 16 kHz, 24 kHz and 32 kHz) at various sound levels (from 0 to 90 dB) ranging to cover the mouse auditory frequency range. ABR measures of each animal were performed individually and using OtoPhyLab system. Evoked potentials were extracted by the signal averaging technique foreach noise level and ABR thresholds for each frequency were determined using OtoPhyLab software. Only the left ears were analyzed in this study.

### Electroretinogram ERG

Mice were anesthetized using ketamine/xylazine mixture. Both eyes were treated with 1% atropine sulfate, 2.5% phenylephrine hydrochloride and 0.5% proparacaine hydrochloride, allowing drops to sit on the eyes for ~2 min before wicking with a cotton swab and applying the next drop. Then, the mouse was positioned on the heated platform. The ground needle electrode was placed in the base of the tail, reference needle electrode subdermally between the eyes and the contact lens electrodes onto the corneas. The scotopic (dark-adapted) ERG measures were performed at 0.001 cd s/m^2^; 0.1cd s/m^2^; 1 cd s/m^2^; and 10 cd s/m^2^ with 5 trials by step, pulse frequency: 0.2 Hz, sample frequency: 1.000 Hz, trial pre-trigger time: 20 msecand trial post-trigger time: 250 msec. At the end of the measure, electrodes were removed from the mouse and the animal was placed in a clean cage on top of a heat pad until recovering.

### Statistical analysis

Descriptive statistics by groups were expressed as mean ± SEM for continuous variables. Statistical significances were determined using a 2-tailed Student’s t test or 2-way ANOVA, followed by a Bonferroni multiple comparisons post hoc test, allowing comparisons between groups versus control, assuming the normal distribution of the variable and the variance homoscedasticity. Statistical analyses were performed using GraphPad Prism version 5.02 for Windows, GraphPad Software, La Jolla California USA.A P value of less than 0.05 was considered significant. n= 5 animals/group.

## Results

### Neuromuscular impairment in CMT mouse at 6 months old

Three different behavioral tests were used to evaluate the neuromuscular impairment of CMT mice: cross time of the balance beam to evaluate walking performance, rotarod latency to study coordination and equilibrium, and Kondziela’ inverted screen test to determine neuromuscular strength.

Similar balance beam cross times were observed in control and CMT groups at the baseline (1 month old). However, the beam cross time was increased at 15.3 ± 1.7 seconds in the CMT group mice whereas the cross time of control animals remained at 9.0 ± 1.0 seconds (Figure 1A)

**Figure 1.**
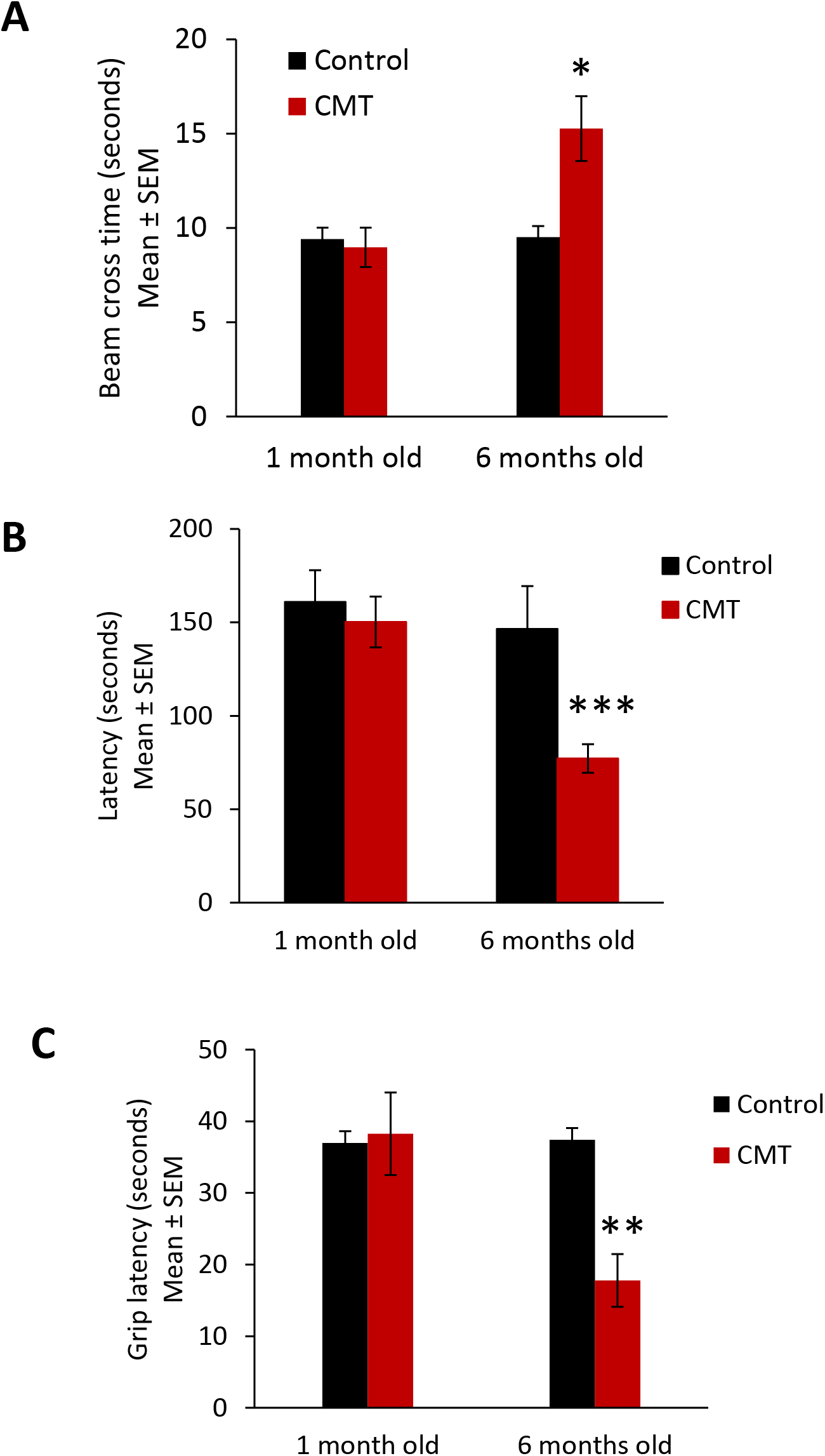
Neuromuscular impairment observed in CMT mice. (A) Balance beam cross time, (B) rotarod latency, and (C) Kondziela’ inverted screen latency of control (black bar) and CMT mice (red bar) at the baseline (1 month old) and at 6 months old. Error bars indicate SEM. Statistical tests are repeated measures two-way ANOVA test comparing CMT to control mouse values. * p<0.05, ** p<0.01, *** p<0.001, no significant (ns).

Using roratod test, we observed similar latency in control and CMT groups at baseline (1 month old). But a significant decrease of rotarod latency was observed in the CMT group compared to the control group at 6 months old. The CMT group presented a mean of rotarod latency of 77.1 ± 7.7 seconds at 6 months old whereas the latency of control animals remained at 150.2 ± 13.6 seconds (Figure 1B).

Finally, similar grip latencies were observed in control and CMT groups at 1 month old. However, at 6 months old, the grip latency significantly decreased at 17.8 ± 3.7 seconds in the CMT group mice whereas the latency of control animals remained at 38.3 ± 5.8 seconds (Figure 1C).

Taken together, these results confirm the presence of a degenerative neuromuscular phenotype in the CMT strain mouse at 6 months old.

### Sciatic nerve electrophysiology impairment in CMT mouse at 6 months old

Behavioral data was confirmed using electromyography of control and CMT sciatic nerve at 1 month old and 6 months old.

Concerning the compound muscle action potential (CMAP) amplitude no significant differences were observed in amplitude between control and CMT at the baseline and at 6 months old (Figure 2A). However, when we determined the nerve conduction velocity (NCV), a significant decrease of the nerve conduction velocity at 8.3.0 ± 2.7 m/s was observed in the CMT group mice whereas the velocity of control animals was 40.5 ± 2.5 m/s (Figure 1B). The action potential velocities of sciatic nerve of control and CMT were similar at 1 month old.

**Figure 2.**
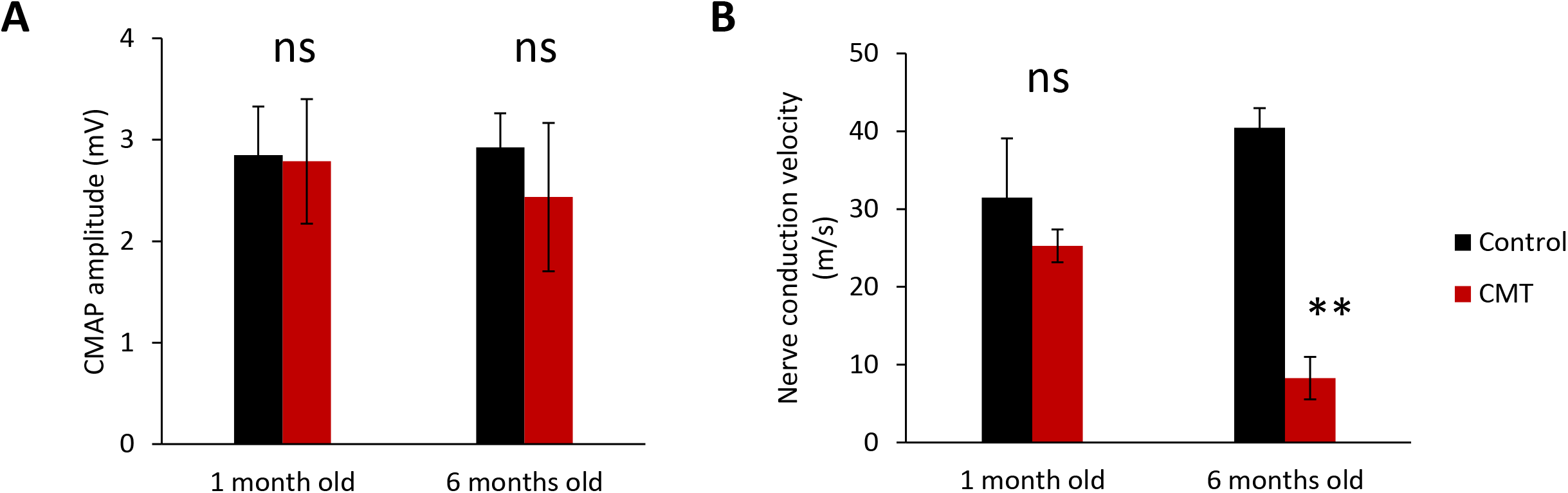
Sciatic nerve electrophysiology impairment observed in CMT mice. (A) Compound muscle action potential amplitude and (B) nerve conduction velocity of sciatic nerve of control (black bar) and CMT mice (red bar) at the baseline (1 month old) and at 6 months old. Error bars indicate SEM. Statistical tests are repeated measures two-way ANOVA test comparing CMT to control mouse values. * p<0.05, ** p<0.01, *** p<0.001, no significant (ns).

### Hearing loss and visual impairment observed in CMT mouse at 6 months old

Peripheral nerve electrophysiological impairment usually leads to sensory neuropathies. For this reason, we determine the hearing and visual performances of control and CMT mice at the baseline (1 month old) and at 6 months old.

Hearing impairment was observed in CMT mice at 6 months old. ABR thresholds of the control and CMT groups were not significantly different at the baseline (1 month old) (Figure 3A).

**Figure 3.**
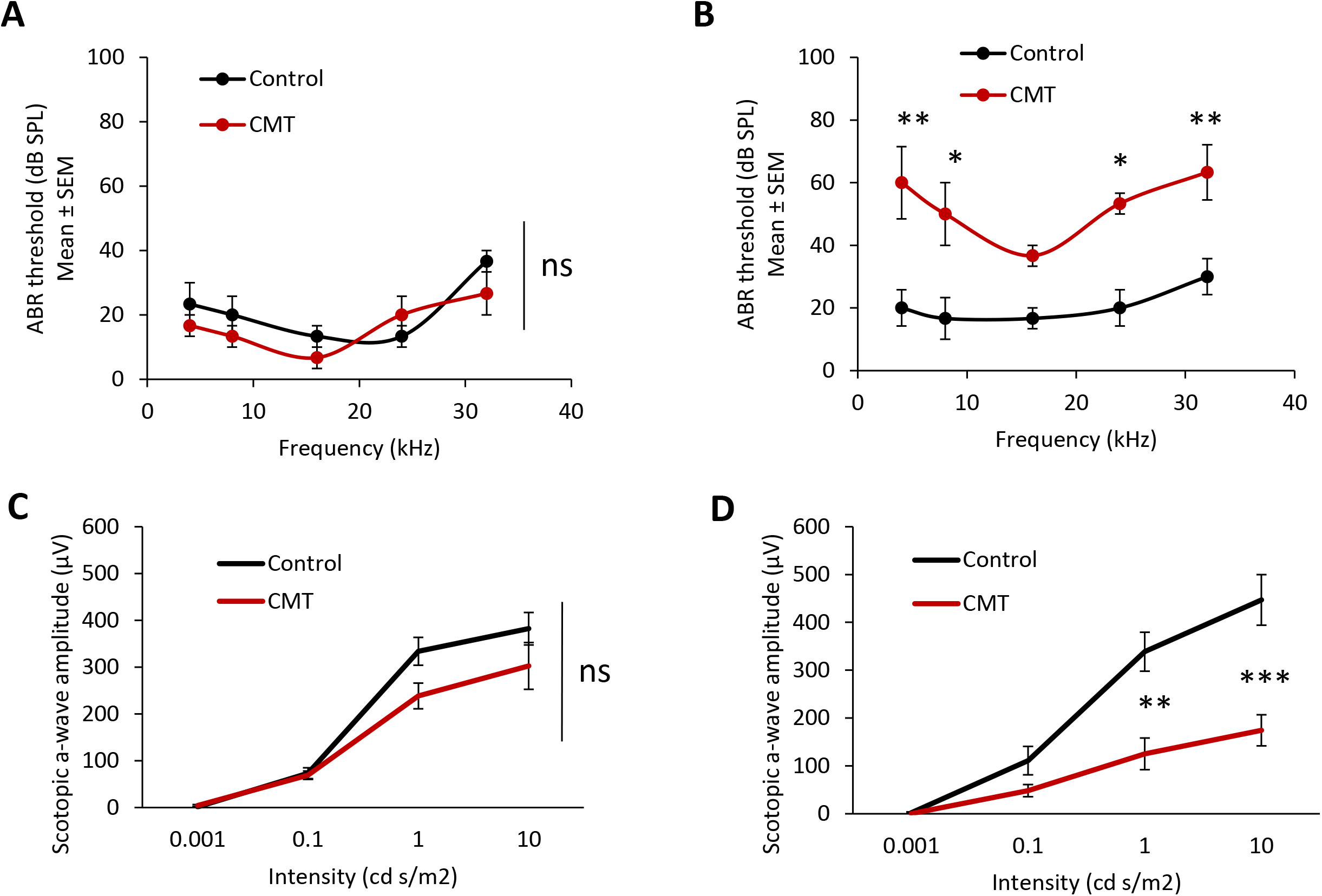
Hearing loss and visual impairment observed in CMT mice. Auditory brainstem responses of control (black line) and CMT mice (red line) at 1 month old (A) and 6 months old (B). Electroretinogram of control (black line) and CMT mice (red line) at 1 month old (C) and 6 months old (D). Error bars indicate SEM. Statistical tests are repeated measures two-way ANOVA test comparing CMT to control mouse values. * p<0.05, ** p<0.01, *** p<0.001, no significant (ns).

No significant differences in ABR thresholds were observed between 1 month old and 6 months old in the control group. However, in the CMT group, the ABR thresholds significantly increased at 4 kHz, 8 kHz, 16 kHz, 24 kHz and 32 kHz at 6 months old compared to the baseline (1 month old). Moreover, the ABR thresholds of CMT mice significantly increased at 4 kHz, 8 kHz, 24 kHz and 32 kHz at 6 months compared to the control group (Figure 3B).

Electoretinogram (ERG) was performed to determine visual performance. Retina ERG impairment was observed in CMT mice at 6 months old. Scotopic a-wave amplitude of the control and CMT groups were not significantly different at the baseline (1 month old) at the analyzed intensities (Figure 3 C).

No significant differences in scotopic a-wave amplitudes were observed between baseline and 6 months old in the control group. However, in the CMT group, the scotopic a-wave amplitude significantly decreased at 1 cd s/m^2^ and 10 cd s/m^2^ at 6 months old compared to the baseline. In the same way, the scotopic a-wave amplitudes of CMT mice were significantly lower at 1 cd s/m^2^ and 10 cd s/m^2^ at 6 months compared to the control group (Figure 3D)

### Myelin and axonal degeneration in CMT mouse at 6 months old

Histological analysis of sciatic nerve and auditory nerve were perfomed in CMT and control mice at 6 months old. Strong Schwann cell demyelination, remyelinating phenotype (after demyelination) and reduction of the myelin sheath were observed in sciatic nerve of CMT mice at 6 months old compared to control mice (Figure 4A). In auditory nerve, as observed in sciatic nerve, strong Schwann cell demyelination and disorganization of the myelin sheath were observed in CMT mice at 6 months old compared to control mice (Figure 4B).

**Figure 4.**
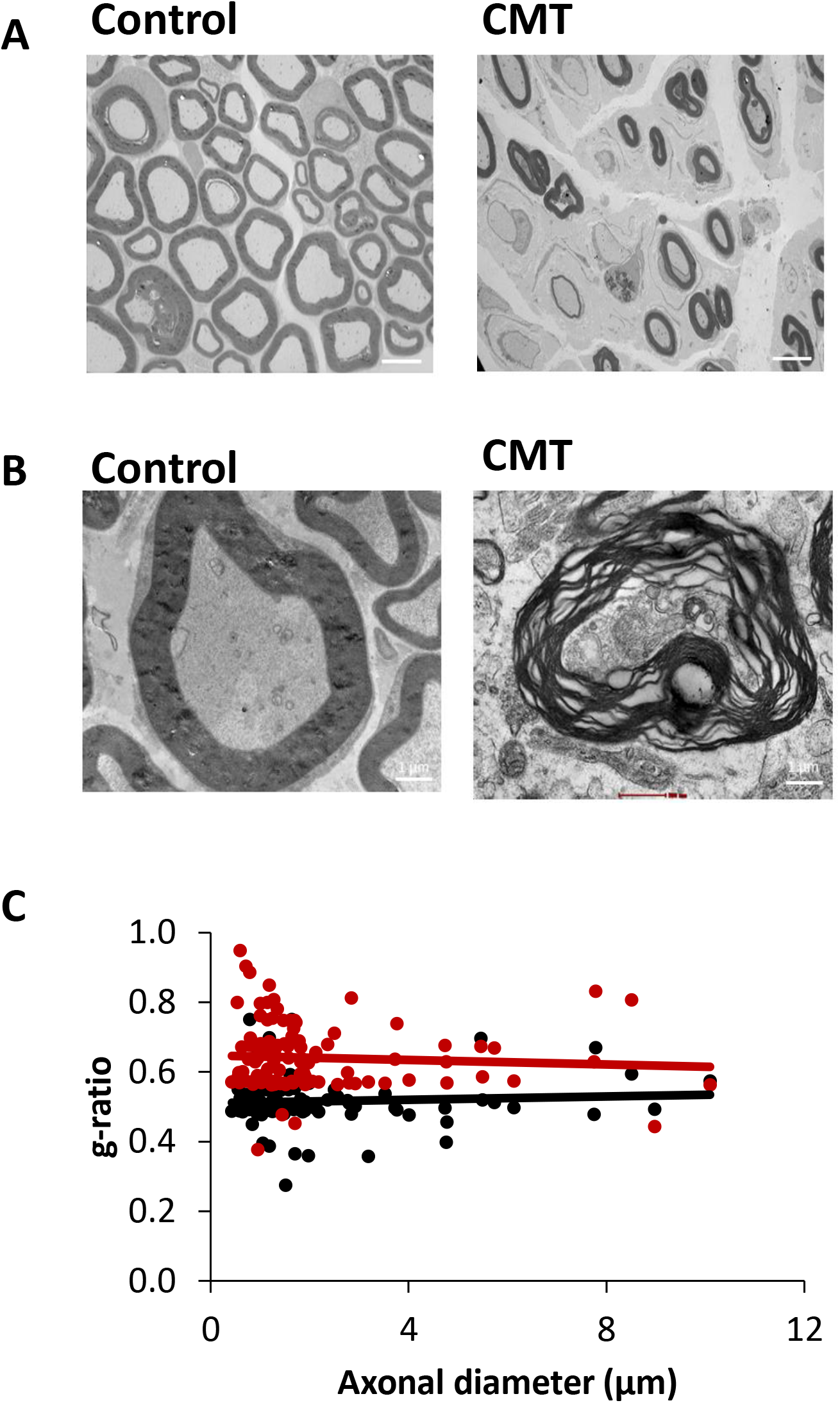
Histological analysis of sciatic and auditory nerves. (A) Transmission electron microscopy images of sciatic nerve of control and CMT mice at 6 months old. Scale bar : 4 μm. (B) Transmission electron microscopy images of auditory nerve of control and CMT mice at 6 months old. Scale bar : 1 μm. (C) Quantification of g ration (myelin sheath) of sciatic nerve of control (black line) and CMT mice (red line) at 6 months old.

In sciatic nerve, the reduction of myelin sheath leaded to a significant increase of g-ration at 0.6408 ± 0.009 for CMT mice whereas the control mice presented a g-ratio mean of 0.515 ± 0.0062 (Figure 4C)

## Conclusion

Decrease of sciatic nerve action potential velocity, neuromuscular disorders, sciatic nerve and auditory nerve demyelination, hearing loss and decrease of scotopic a-wave amplitude (visual acuity) was observed in CMT1A mice at 6 months old.

Taken together, these results confirm that the C3-PMP22 mouse is a robust and reproducible model to analyze the sensory neuropathy and neuromuscular disorders observed in CMT 1A disease. This model allows to determine the efficacy of new pharmacological candidates targeting CMT1A disorder.

## Notes

### Competing Interest Statement

The authors have declared no competing interest.

